# A proteome-wide yeast degron collection for the dynamic study of protein function

**DOI:** 10.1101/2024.06.10.598194

**Authors:** Rosario Valenti, Yotam David, Dunya Edilbi, Benjamin Dubreuil, Yeynit Asraf, Tomer Meir-Salame, Ehud Sass, Maya Schuldiner

**Affiliations:** Weizmann Institute of Science

**Keywords:** protein depletion, degron, yeast genomics, yeast mutant array, cell cycle, mitochondrial morphology

## Abstract

Genome-wide collections of yeast strains, known as libraries, revolutionized the way systematic studies are carried out. Specifically, libraries that involve a cellular perturbation, such as the deletion collection, have facilitated key biological discoveries. However, short-term rewiring and long-term accumulation of suppressor mutations often obscure the functional consequences of such perturbations. We present the AID library which supplies “on demand” protein depletion to overcome these limitations. Here, each protein is tagged with a Green Fluorescent Protein (GFP) and an Auxin inducible degron (AID), enabling rapid protein depletion that can be quantified systematically using the GFP element. We characterized the degradation response of all strains and demonstrated its utility by revisiting seminal yeast screens for genes involved in cell cycle progression as well as mitochondrial distribution and morphology. In addition to recapitulating known phenotypes, we also uncovered proteins with previously unrecognized roles in these central processes. Hence, our tool expands our knowledge of cellular biology and physiology by enabling access to phenotypes that are central to cellular physiology and therefore rapidly equilibrated.

## Introduction

The baker’s yeast, *Saccharomyces cerevisiae*, stands as a cornerstone in the exploration of cellular biology due to its genetic tractability, cost-effectiveness, and substantial evolutionary conservation with humans (Cohen, Kahana, and Schuldiner 2022; Laurent et al. 2020). The advent of genomic collections, also known as libraries, has been a transformative force in yeast studies, enabling genome-wide investigations through arrayed ensembles of strains, each modified at a single gene to facilitate comprehensive studies (Botstein and Fink 2011).

Notably, the pioneering deletion library (Yeast Knock-Out or YKO) (Giaever et al. 2002; Winzeler et al. 1999), where over 90% of the non-essential genes were deleted in haploid strains, has played a fundamental role in numerous key biological discoveries, accumulating thousands of citations (Giaever and Nislow 2014). Systematic studies based on the KO library allowed the association of genes with a specific phenotype and from that, cellular roles. For example, the screen for genes required for correct mitochondria distribution and morphology (MDM), carried out by staining the mitochondria of the YKO collection and identifying strains with altered mitochondria, enabled the definition of MDM strains – many of which were shown to be directly involved in mitochondrial shaping and inheritance (Dimmer et al. 2002). Complementary to the YKO library, the Green Fluorescent Protein (GFP) library (Huh et al. 2003) established a landmark in the systematic visualization of over 60% of the GFP-fused proteome, providing valuable insights into protein abundance and subcellular localization.

However, the construction of such libraries was historically time-consuming and expensive until the development of the SWAp-Tag (SWAT) libraries (Meurer et al. 2018; Weill et al. 2018; Yofe et al. 2016). The SWAT libraries harbor a placeholder modification in the amino-(N’) or carboxy-(C’) terminus for easy substitution by any genetic sequence of choice such as a modification (tag), promoter, 5’ untranslated region (UTR), 3’ UTR or selection cassette. Despite these advancements, approximately a fifth of yeast genes remain uncharacterized or poorly characterized (Cohen et al. 2022; Gaikani et al. 2024). Partially, this can be attributed to limitations in existing tools, such as compensatory mutations arising after gene deletion (Hughes et al. 2000; Teng et al. 2013) that would mask informative phenotypes, and short-term cellular rewiring, that obscures the correct coupling between perturbation and observed phenotype.

Addressing these challenges, conditional libraries aimed at altering gene product abundance were developed. For example, Tetracycline-controlled gene expression from the TET-off system (Mnaimneh et al. 2004), can repress transcription of the tagged gene upon addition of doxycycline, a more stable analogue of the tetracycline antibiotic. Alternatively, estradiol allows the induction of translation in yeast strains from the YETI library (Yeast with Estradiol strains with Titratable Induction) (Arita et al. 2021). Not surprisingly, essential genes are the main focus of these conditional libraries since, by definition, they were absent from the YKO library. Although useful, these approaches met partial success in eliciting expected phenotypes, potentially due to residual cellular rewiring. Perturbing the translation of a gene, or even the levels of its mRNA (Schuldiner et al. 2005), might be too slow to report on an immediate phenotype, as seen when contrasting phenotypes of the heat-inducible-degron (Kanemaki et al. 2003) with literature reports on phenotypes visualized by other methods (Yu et al. 2006). Indeed, rapid protein destabilization, as the one achieved with thermosensitive (TS) strains (Kofoed et al. 2015; Li et al. 2011) has been historically used to probe the role of essential components and was instrumental in the discovery of the Cell Division Cycle (CDC) genes (Hartwell et al. 1973; Hartwell, Culotti, and Reid 1970). CDC mutants in TS strains grown in their restrictive temperature perished while displaying a synchronized morphology, setting the basis for the discovery of the cyclins and cyclin-dependent kinases currently known as governing the cell cycle progression. This example not only highlights the power of yeast genetics, but also the complementarity that different strategies offer and the importance for having tools for rapid, on demand, protein depletion.

Motivated by the notion that new tools drive biological advancements, we opted for a degron-based conditional protein depletion strategy. We chose the improved Auxin Inducible Degron (AID2) system (Yesbolatova et al. 2020) as a rapid, reversible and flexible system that relies on the addition of a small molecule, a chemically modified version of auxin, 5-Phenyl 1H-indole-3-acetic acid (5-Ph-IAA), for its induction of protein depletion. This system enjoys baseline residual degradation in the absence of external stimulus thus making it much less taxing on cells and more reactive to experimental conditions. Leveraging the C’ SWAT library, we extended our approach to the entire proteome allowing us to screen for immediate phenotypes following degradation of both non-essential and essential proteins. We also incorporated in our design a GFP tag for simultaneous reporting on protein abundance, localization, and the effects of protein deletion — a conceptual fusion of the GFP library with the YKO.

We validated our C’ AID-GFP library by revisiting seminal yeast studies, uncovering additional essential proteins under different growth conditions, recapitulating and extending the MDM screen, and systematically detecting strains with synchronized morphologies, in a re-approach of the CDC screen. Successful reproduction of these studies and identification of new genes functions attests the complementarity of our approach within the expansive toolbox of yeast genetics. Our library will be freely distributed hoping to fuel a wave of discoveries on yet uncharacterized yeast protein functions.

## Results

### Establishing a system for fast and tractable protein depletion based on the AID2 system

To induce physiologically relevant and informative perturbations while minimizing cellular rewiring, our goal was to establish a rapid and efficient protein depletion system. To this aim, we adapted the improved Auxin Inducible Degron system (AID2), a versatile tool for dynamic, on-demand depletion of endogenous proteins in yeast cells (Yesbolatova et al. 2020). This system is superior to previous ones in that background degradation by auxin-like cellular molecules is dramatically diminished, minimizing background degradation in absence of the inducer. The system is also more specific, requiring a low dose of inducer that presents no toxicity to the cells and that can be applied under diverse conditions without generating an inherent stress to the cells.

Our implementation of the AID2 system involves fusing the small AID* tag (amino acids 71-114 from AtIAA17, Uniprot ID:P93830, from hereon termed simply AID) (Morawska and Ulrich 2013) followed by the enhanced, eGFP (Zacharias et al. 2002), for visualization, to the C’ of a protein, on the background of constitutive expression of the adaptor OsTIR1(F74G). The system is induced by addition of the modified auxin, 5-Ph-IAA, which leads to the recognition of the AID tag on the protein of interest by the TIR1 adaptor protein. TIR1 binding to the AID recruits the ubiquitination machinery (Skp1-Cul1 E3 ligases (Yesbolatova et al. 2020; Yu et al. 2015)) and leads to proteasomal degradation of the AID-tagged protein (Figure 1A).

**Figure 1.**
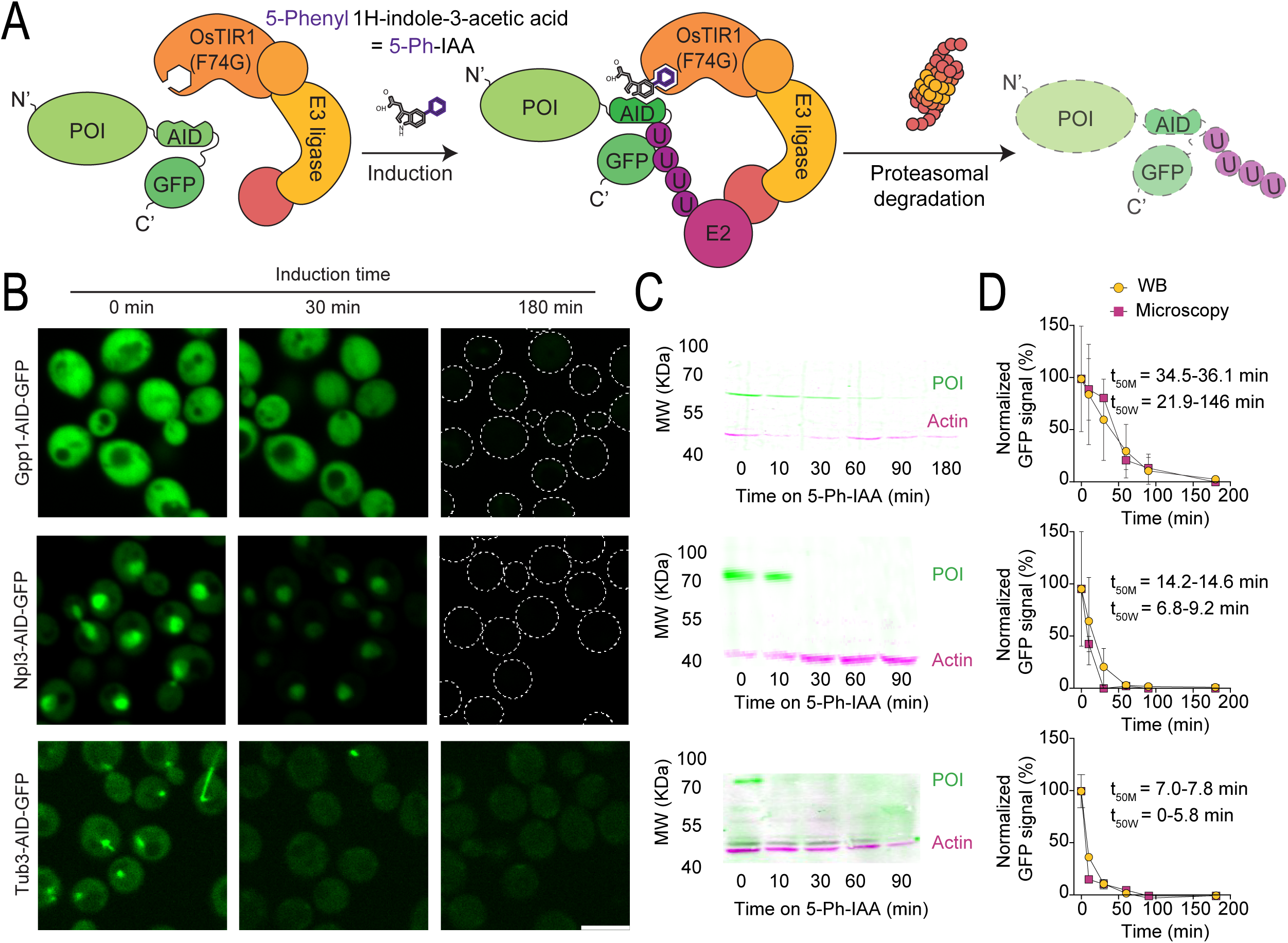
The Auxin Inducible Degron (AID) system allows “on demand” depletion of the proteome. A) Schematic representation of the AID system, where a <u>p</u>rotein <u>o</u>f interest (POI) is fused to the AID2 tag and to a GFP for visualization. In the same cells, the OsTIR1(F74G) adaptor protein is constitutively expressed. Upon induction by the addition of the auxin analog 5-PH-IAA, the construct gets ubiquitinated and degraded via the proteasome. B) Fluorescent images of selected proteins tagged with the AID system after 0, 30 and 180 minutes of induction. All the proteins are completely or partially depleted upon induction. For Gpp1 and Npl3 at 180min, the outline of the cells is depicted with a dotted line. Scale bar: 5µm. C) Western blots (WB) of the same strains shown in B. Immunoblotting was performed against GFP (green) and actin (magenta) as a loading control. The GFP-tagged proteins are completely or partially depleted upon induction. Only one band is detected, at the expected molecular weight in each case, indicating no free GFP or intermediate degradation stages are generated. D) Graphs comparing the degradation curves of each protein from B, as determined based on the fluorescence images in B (yellow circles) or on the western blots in C (pink squares), normalized to time 0. Both methods produced comparable curves, despite the half-life (t_50M_ for microscopy and t_50W_ for WB) of the proteins spanning a wide range of times. Three technical replicates were used in each case. A one-phase decay curve was fitted to the data to estimate the half-lives of each protein and the 95% confidence interval of the t_50_ is displayed.

To ensure that GFP fluorescence accurately reports on protein degradation rates, we monitored the degradation of various proteins using both GFP fluorescence imaging (Figure 1B) as well as western blot analysis (Figure 1C). We confirmed the correspondence between protein levels and GFP fluorescence (Figure 1D), irrespective of the protein’s half-life, and also that free GFP does not accumulate during degradation. Since GFP imaging presented higher throughput and additional insights into the subcellular localization of the remaining protein, we turned to measuring GFP fluorescence changes as a reliable indicator of protein degradation.

### Creating a proteome-wide collection of depletable proteins and characterizing responsiveness at a proteome level

To enable comprehensive characterization of yeast phenotypes following “on demand” elimination, we extended the modified AID2 system to a whole-proteome yeast library using the recently developed SWAT approach for library generation (Meurer et al. 2018; Weill et al. 2018; Yofe et al. 2016) (Figure 2A). Following the “SWATting” procedure, we had 91% of the strains present in the new library (survival rate (Supplementary Figure 1A)). We estimated 94% of “SWATting” efficiency, considering the fraction of strains with effectively detected fluorescence signal among the 500 most abundant proteins (Ho, Baryshnikova, and Brown 2018) (Supplementary Figure 1B).

**Figure 2.**
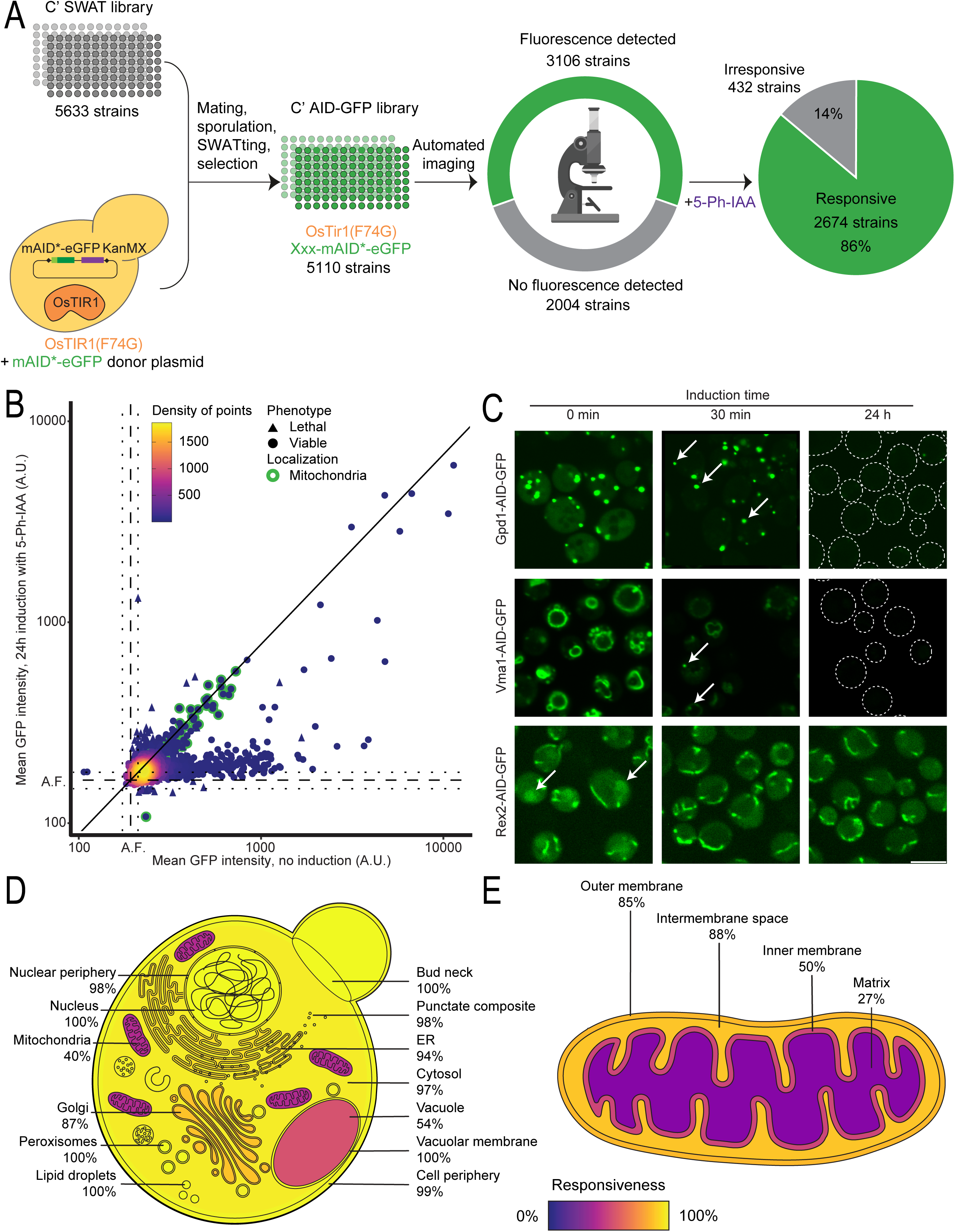
The C’ AID-GFP library has robust responsiveness across cellular compartments. A) Summary schematics of the expansion of the AID system to a whole genome collection. The original C′ SWAT library was used to generate the C’ AID-GFP library using an automated process of mating, selection, sporulation and SWATing. In the final library, every strain expresses the OsTIR1(F74G) protein, and a different gene is fused to the AID-GFP tag. Fluorescence was detected in 60% of the strains under standard laboratory conditions (log-phase cells in rich media). Of those, 86% responded to the induction with a measurable protein depletion. B) Dot plot showing the mean GFP fluorescence intensity per cell for each strain before or after 24h induction time. The background autofluorescence noted A.F. is calculated as the mean fluorescence of control strains is marked in dotted lines with a confidence interval of two standard deviations in each direction. The diagonal line indicates the expected location for irresponsive strains. Many strains die after induction (triangles), making their fluorescence signal not informative, while most of the irresponsive strains encode for mitochondrial proteins (green edge). C) Fluorescent images of selected proteins imaged after no (0 min), short (30 min) or long (24h) periods of induction. In the case of dually localized proteins, induction depletes one of the subpopulations with faster kinetics. Gdp1 remains visible in peroxisomes, Vma1 in endosomes and Rex2 in mitochondria while their respective cytosolic, vacuolar, or nuclear subpopulations are depleted. In cases where the signal is too dim, the outline of the cells is depicted with a dotted line. Scale bar 5µm. D and E) Schematics of a yeast cell (D) and a mitochondrion (E) displaying the responsiveness per subcellular compartment. Proteins in membrane-bound organelles are more protected from degradation but the responsiveness across sub-cellular compartments is still high.

We also sequenced most of the library to validate “in frame” insertion of the tag, that was confirmed for 3293 strains (Supplementary Figure 1C and D; Supplementary Table 1). Hence, we successfully created the C’ AID-GFP library comprised of 5,110 arrayed strains, each constitutively expressing the TIR1 adaptor protein and containing a unique gene fused with the AID tag and GFP at their C’ terminus.

Upon imaging the entire collection, approximately 60% of the strains exhibited a fluorescent signal, consistent with reports suggesting that nearly 2,000 proteins are either not expressed at all or are very low abundance under standard laboratory conditions (log-phase growth in rich media) (Breker, Gymrek, and Schuldiner 2013; Ghaemmaghami et al. 2003; Huh et al. 2003). Inducing the AID system led to the complete or partial degradation of nearly 90% of the strains evaluated under these conditions (Figure 2A). This high responsiveness underscores the broad applicability of our system for effectively knocking down proteins in diverse cellular contexts.

Since we observed variations in the depletion dynamics among individual proteins (Figure 1B-D), we systematically analyzed the response of each strain to short (30 min) and long (24h) periods of induction. We imaged the AID strains by high-throughput fluorescence microscopy before and after induction, with actively dividing cells in rich media. The resulting images were manually inspected, and we determined responsive strains with significant reduction of their GFP signal after induction combining the manual examination results with a comprehensive quantitative analysis. Within our collection, we identified proteins that exhibited rapid depletion, such as Lia1, while others required extended induction time, exemplified by Mrh1. Concurrently, some proteins, like Tim44, displayed negligible responsivity to the AID system even after a long induction time of 24h (Supplementary Figure 2A).

A short induction time was sufficient to measure changes for 1642 strains (Supplementary Figure 2B), although a clearer response emerged after 24 hours of induction, with 2674 strains, representing 86% of the visible proteome, displaying responsivity (Figure 2B). Notably, after this extended induction, we observed the demise of certain strains (Figure 2B, triangles). This induction-provoked lethality implies the depletion of an essential protein in those strains, confirming that the AID-tagged protein was crucial for cell viability. If we take these strains too into consideration than overall our library has 2713 clearly responsive strains.

Interestingly we noticed that over 100 dually localized proteins exhibited distinct depletion kinetics for each subcellular population (Figure 2C). Exploiting these differential degradation rates provided enhanced visualization of the eclipsed subpopulations, particularly notable after short induction periods (Figure 2C).This enables visualizing localizations for several proteins previously thought to be solely cytosolic.

Additionally, we expected proteins behind cellular membranes to face challenges accessing the ubiquitination targeting and proteasomal degradation machinery upon induction. Indeed, we observed many irresponsive strains with their tagged protein localized to the mitochondria (Figure 2B, green edge) or to the lumen of other organelles in line with our expectations. Despite this limitation, the responsiveness per cellular compartment was robust, ranging from 40-100% (Figure 2D). Even within mitochondria, responsiveness correlates with the sub-organellar compartment, with the mitochondrial matrix proving to be the most challenging sub-compartment to target (Figure 2E). We attribute the weaker response to the time spent by proteins dwelling in the cytosol before translocating into mitochondria, though milder depletion could also reflect longer protein half-lives (Bomba-Warczak and Savas 2022). Proteins lingering longer in the cytosol are more susceptible to the clearance of newly synthesized proteins. The remaining protected protein pool may be diluted by cellular divisions or depleted through natural turnover. Segregating mitochondrial proteins into co-and post-translationally translocated groups (Williams, Jan, and Weissman 2014) (Supplementary Figure 2C) supports our hypothesis about higher degradation with increased residency in the cytosol. Notably, post-translationally translocated proteins are 2.2 times more likely to be responsive to induction of degradation. A similar trend is observed with the C’ orientation of membrane proteins (Weill et al. 2019), where exposure of the AID element to the cytosol, leads to more efficient degradation, as exemplified for the Endoplasmic Reticulum (ER) (Supplementary Figure 2D).

Overall, even when protein abundance correlated with quantified fluorescence as expected, (Supplementary Figure 2E), the responsiveness of the fluorescent fraction remained quite steady (Supplementary Figure 2F). Between the first and last decile of abundance, responsiveness ranged from 82 to 92% ruling out the dependence of induced degradation on protein abundance. This seems to indicate that the ubiquitin proteasome system is not a limiting factor and therefore responsiveness is not conditioned by the abundance of the protein.

### Application of the C’ AID-GFP library to the study of essential proteins

To assess the utility of our library, we revisited seminal yeast studies focusing on central cellular processes and examined our collection’s ability to recapitulate known phenotypes as well as its potential to uncover genes not previously known to affect these processes.

Encouraged by the observation of induction-provoked lethality in certain strains (Figure 2B), our initial investigation concentrated on determining protein essentiality. To this end, we cultured our arrayed yeast collection on three types of solid media that represent different yeast growth physiologies: rich media, minimal media or a non-fermentable carbon source; with or without 5-Ph-IAA, to induce the degradation of the tagged proteins. Colony sizes were measured after growth in induced versus uninduced conditions. The ratios of colony sizes scored the extent of the relative growth defect due to induced protein degradation among the strains (Figure 3A). Most strains grew similarly across different media including 347 strains presenting severe defects (characterized by a relative growth score below 0.5) across all three media tested (Figure 3B and Supplementary table 1). In addition, 254 strains manifested growth defects in only one or two of the tested media (Figure 3C).

**Figure 3.**
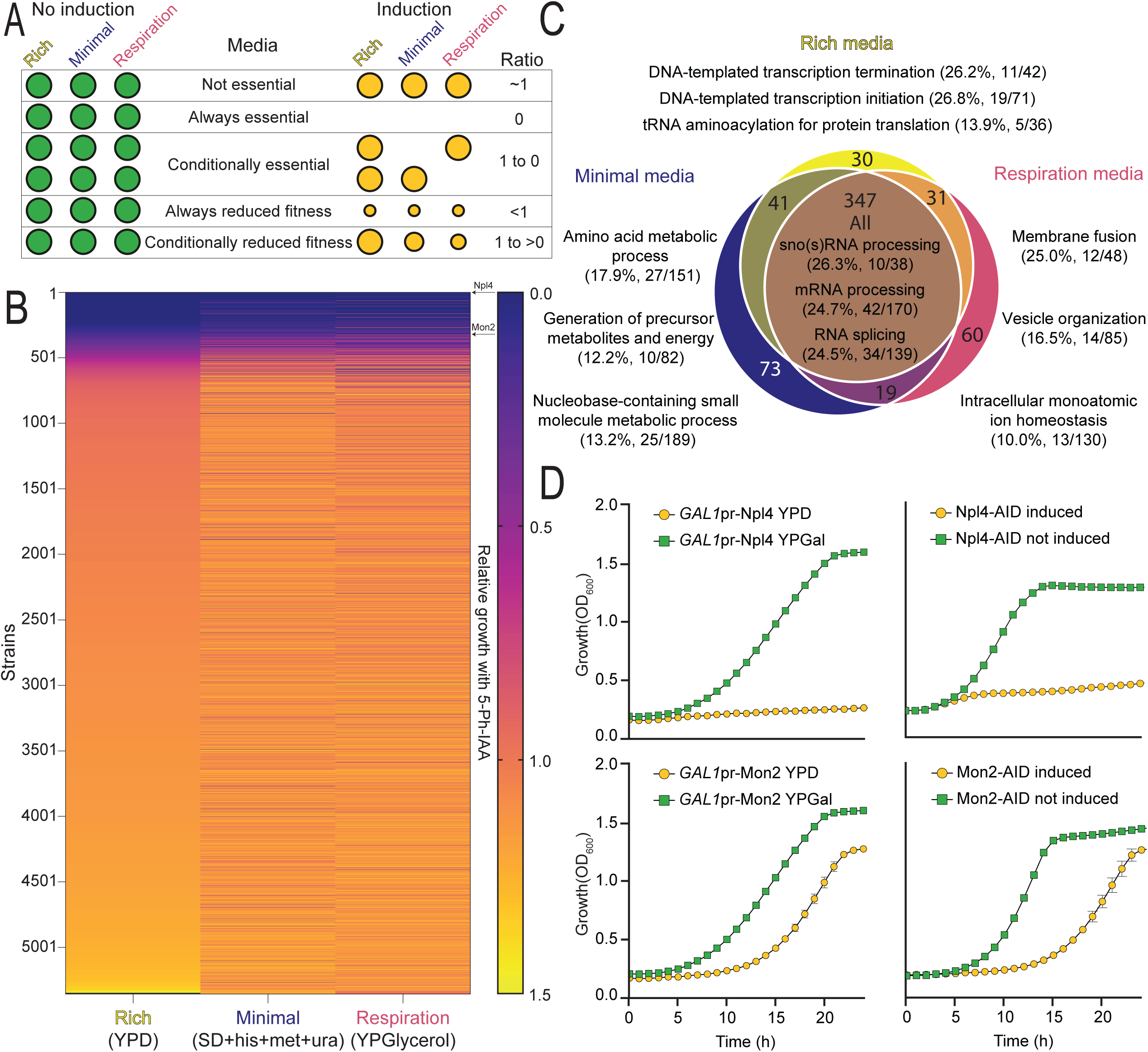
The C’ AID-GFP library can be applied to the study of protein essentiality. A) Schematics representing our pipeline to test protein essentiality in different media. The arrayed yeast collection can be grown in different types of solid media with and without induction of the AID system. The ratio of the colony size, or relative growth, is indicative of whether the protein depleted is important for growth rate or viability. B) Heatmap showing the relative growth of every strain in the collection in three different media. Most strains, but around 500, display no severe growth defect upon induction of degradation. The results are very similar for all the medias tested, with specific strains displaying altered growth in one or more media. Strains were grown in duplicates and for two consecutive plate replications. C) Venn diagram summarizing the results in B for all the strains with a growth defect bigger than 50%. Most of these strains (347) displayed a severe growth defect in every media. The top three GO slim terms for biological processes associated with each group of strains is displayed, together with the percentage of the GO slim term covered in the group. A clear relationship between the media analyzed and the processes affected can be appreciated. D) Growth curves of a selected example for a protein that is essential, as Npl4, or cause a fitness loss, as Mon2, upon depletion (orange series). Both our system and a *GAL1*-promotor swap reproduce the results seen in B. An average of three repeats with their standard deviation is shown.

To compare the results obtained in each media, we calculated the enrichment of Gene Ontology (GO) slim terms related to biological processes (Ashburner et al. 2000; The Gene Ontology Consortium et al. 2023). GO slim, the cut-down version of GO terms, provides a broad overview of biological processes associated with a particular set of genes. We grouped strains with severe growth defects for each media (Figure 3C). We listed the top three associated GO slim terms, along with the percentage of each term recovered in their corresponding group of strains. As anticipated, fundamental processes, such as those related to RNA processing, emerged as top hits for strains affected in all tested media. Processes essential for minimal media, relating to the biosynthesis of amino acids or metabolites, were also prominent in their corresponding group, aligning with the absence of external nutrient provision.

Analysis of all strains displaying severe defects also uncovered new candidate essential genes that were previously missed. The first systematic classification of protein essentiality in yeast resulted from the creation of the YKO library, where the definition of an essential protein came from the unattainable gene deletion of haploid strain despite being created as a heterozygous diploid (Giaever et al. 2002). Examining only our responsive strains, the list of strains with severe growth defects (relative growth threshold <0.5) recovered 69% of those originally defined essential genes (Supplementary Figure 3A, left). To favor the retrieval of *bona fide* unidentified essential genes over slow growing strains, we used a stringent threshold of less than 10% relative growth. Among the 281 strains selected, only 11 had not been identified as essential in the YKO (Supplementary Figure 3A, right), even though five were reported as essential genes in the literature from low-throughput experiments supporting our findings (DeHoratius and Silver 1996; Hampton, Gardner, and Rine 1996; Hoffman et al. 2016; Kastenmayer et al. 2006; Li, Li, and Elledge 2005; Piłsyk et al. 2020; Stirling et al. 2011).

To validate the reproducibility of our high throughput screen, we took two strains and repeated the assay in liquid media using both the AID induction and an orthogonal approach of changing their promoter to a repressible one. Replacing the endogenous promoter with the galactose inducible/glucose repressible, *GAL1*, promoter, we depleted desired proteins by growing cells on media with glucose overnight. Both methods in liquid growth assays recapitulate the results obtained from the high-throughput approach (Figure 3D). In the future, it would be worthwhile to find essential genes under a plethora of conditions, which is now possible since the auxin depletion is robust in a wide array of conditions (for examples see Supplementary Figure 3C). Furthermore, many conditional depletion systems have shown to be more efficient with highly expressed genes (Arita et al. 2021). We noted our system follows the opposite trend (Supplementary Figure 3B), making it highly complementary to other approaches.

### The C’ AID-GFP library uncovers additional proteins required for normal mitochondrial distribution and morphology

Given that mitochondria imposed a tough challenge for our approach, we sought to specifically assess the capability of the collection to yield meaningful insights in mitochondrial cell biology. To address this, we turned to a screen that systematically defined genes essential for correct mitochondrial distribution and morphology (MDM) using the deletion collection for non-essential genes (Dimmer et al. 2002) or the TET-off repressible collection for essential ones (Altmann & Westermann, 2005). Mimicking their approach, we employed a mitochondrial dye to observe mitochondria morphology in strains from our collection, following 24 hours of induction of the degron system.

In line with prior discoveries, we successfully recapitulated known MDM phenotypes (Supplementary Figure 4A). Within our responsive strains, we rediscovered 60% of the proteins previously documented to induce mitochondria morphology defects when depleted in the literature. Differences between the studies may stem from our inability to deplete certain mitochondrial proteins to the extent required for the phenotype to arise, or the result of changes that occur only after a strain has lost its mitochondrial DNA. Such process typically takes longer than 24 hours and would therefore only be present in deletion strains. Moreover, we identified over 220 previously undescribed candidates where mitochondrial morphology appeared altered, as revealed in a double-blind manual analysis and quantification of the images (Supplementary Table 1). Examples of these newly described MDM-like strains (Figure 4A) are proteins from pathways like lipid biogenesis, RNA processing, cytoskeleton organization and many more. This underscores the diverse biological processes regulating mitochondrial morphology and emphasizes the invaluable analytical perspective our library provides in studying mitochondria.

**Figure 4.**
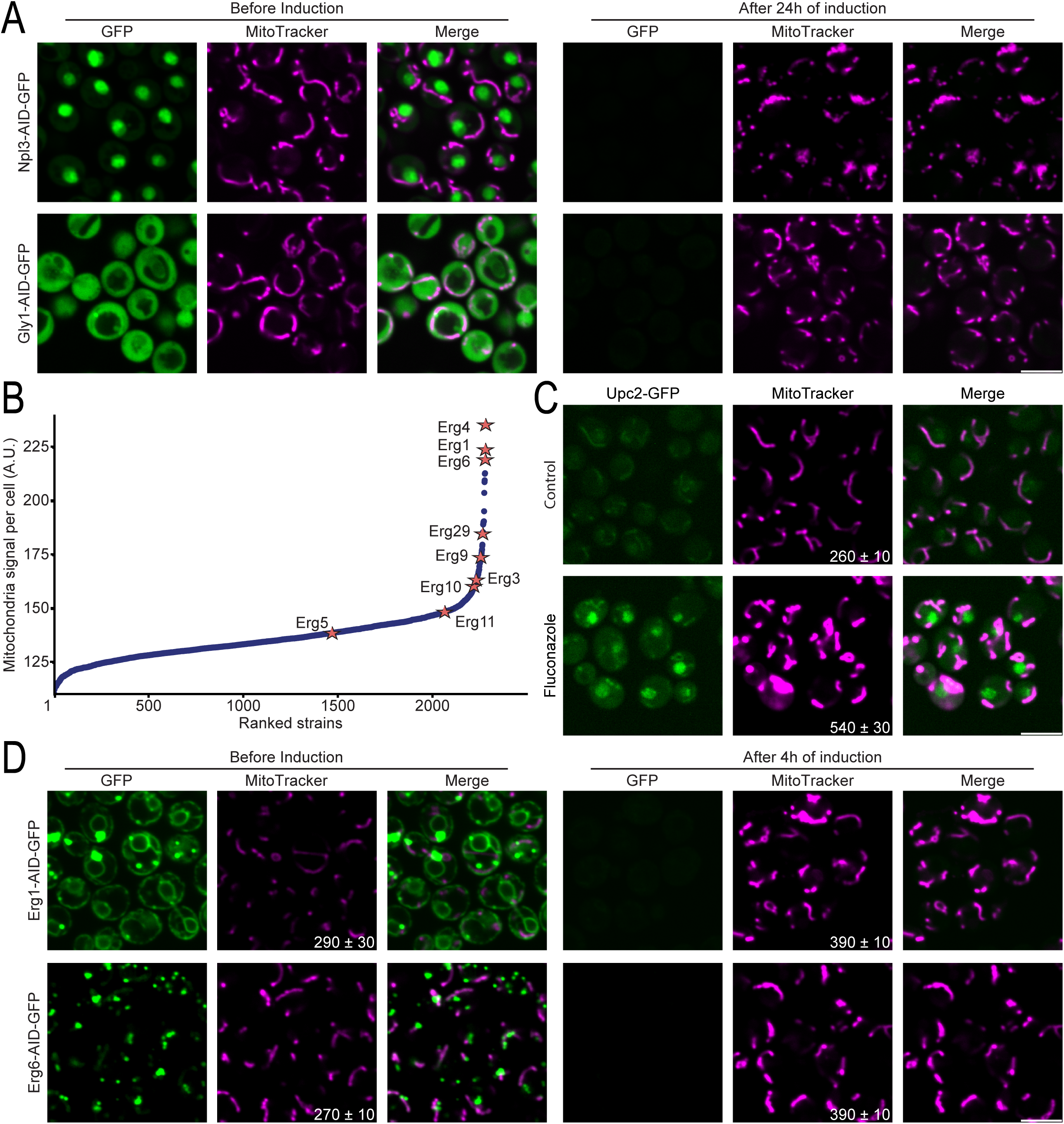
New mitochondrial morphology and distribution phenotypes appear after protein depletion. A) Fluorescent images of strains before and after induction of the degron system for 24h, for previously undescribed proteins required for correct mitochondria morphology and distribution. After depletion of the tagged proteins (green), the mitochondria look aberrant (magenta). B) Scatter plot showing the MitoTracker signal intensity per cell after 24h of induction vs strain rank. Many proteins belonging to the ergosterol biosynthesis pathway (stars) display elevated mitochondrial signal intensity (which could be either changes in membrane potential or in shape) compared to the median. C) Fluorescent images of a strain containing the ergosterol sensitive protein, Upc2, C’ tagged with GFP under control conditions, or upon 4h of treatment with fluconazole, which inhibits ergosterol biosynthesis. Upc2 translocate to the nucleus (green) upon ergosterol depletion. Concomitantly, the MitoTracker signal intensity increases and the mitochondrial morphology changes. D) Fluorescent images of selected strains whose depleted protein participates in the ergosterol biosynthesis pathway, without or with 4h of induction. As seen in C, the mitochondrial morphology changes and the MitoTracker signal intensity increases (magenta) upon protein depletion (green). For all images, scale bar 5µm.

Also in accordance with previous studies (Altmann and Westermann 2005; Zung et al. 2024), systematic evaluation of mitochondria in images after auxin induction revealed increased mitochondrial fluorescence signal owing to the depletion of most ergosterol biosynthesis related proteins (Figure 4B). Since this is an essential pathway, we repeated the experiment but with only 4h of induction of the AID system, to prevent cell death from interfering with our observations. Even after this short depletion time, the mitochondrial morphology and the fluorescent signal’s intensity of most strains were altered (Figure 4D and Supplementary Figure 4B), irrespective of whether they participate in the pre- or post-squalene module of the pathway (Supplementary Figure 4C). Unsurprisingly, exceptions are the strains where Hmg1 or Hmg2 were tagged, in which the depletion of one paralog can be compensated by the action of the other (Basson, Thorsness, and Rine 1986); or Erg11, where the depletion is slower probably due to its C’ facing the lumen of the ER (Supplementary Figure 4B and C). Notably, the morphological modifications were also elicited by fluconazole and terbinafine, two drugs that inhibit the ergosterol biosynthesis pathway (Figure 4C and Supplementary Figure 4C) (Zung et al. 2024). Together our data highlight an important role for sterols in maintain mitochondrial shape and more broadly suggest that additional factors influencing mitochondrial biogenesis await discovery.

### Revisiting the Cell Division Cycle screen using the C’ AID-GFP library suggests additional genes required for cell division

Finally, we revisited the CDC screen (Hartwell et al. 1970), which uncovered proteins essential for normal cell division and cell cycle progression. Since the original screens required the creation of TS strains via mutagenesis, while the C’ AID-GFP library encompass most of the proteome, we speculated there might be proteins with CDC like features still unreported. From our long-term induction screen, it became strikingly apparent that many strains, not only those of previously known CDCs, exhibited altered synchronized morphology (Figure 5A and B). After 24 hours, we systematically identified by visual examination all strains arrested with a synchronized morphology (Supplementary Table 1). Among those strains, we retrieved two well-characterized complex members of one of the original CDC mutants, Cdc48 - Npl4 and Ufd1. Cdc48 has been thoroughly characterized as a AAA ATPase that governs protein degradation and its depletion causes the cells to arrest in G2 phase (Supplementary Figure 5A). Despite not previously being characterized as one, we found that Npl4 displays the same CDC characteristics as its interactor Cdc48 (Supplementary Figure 5B).

**Figure 5.**
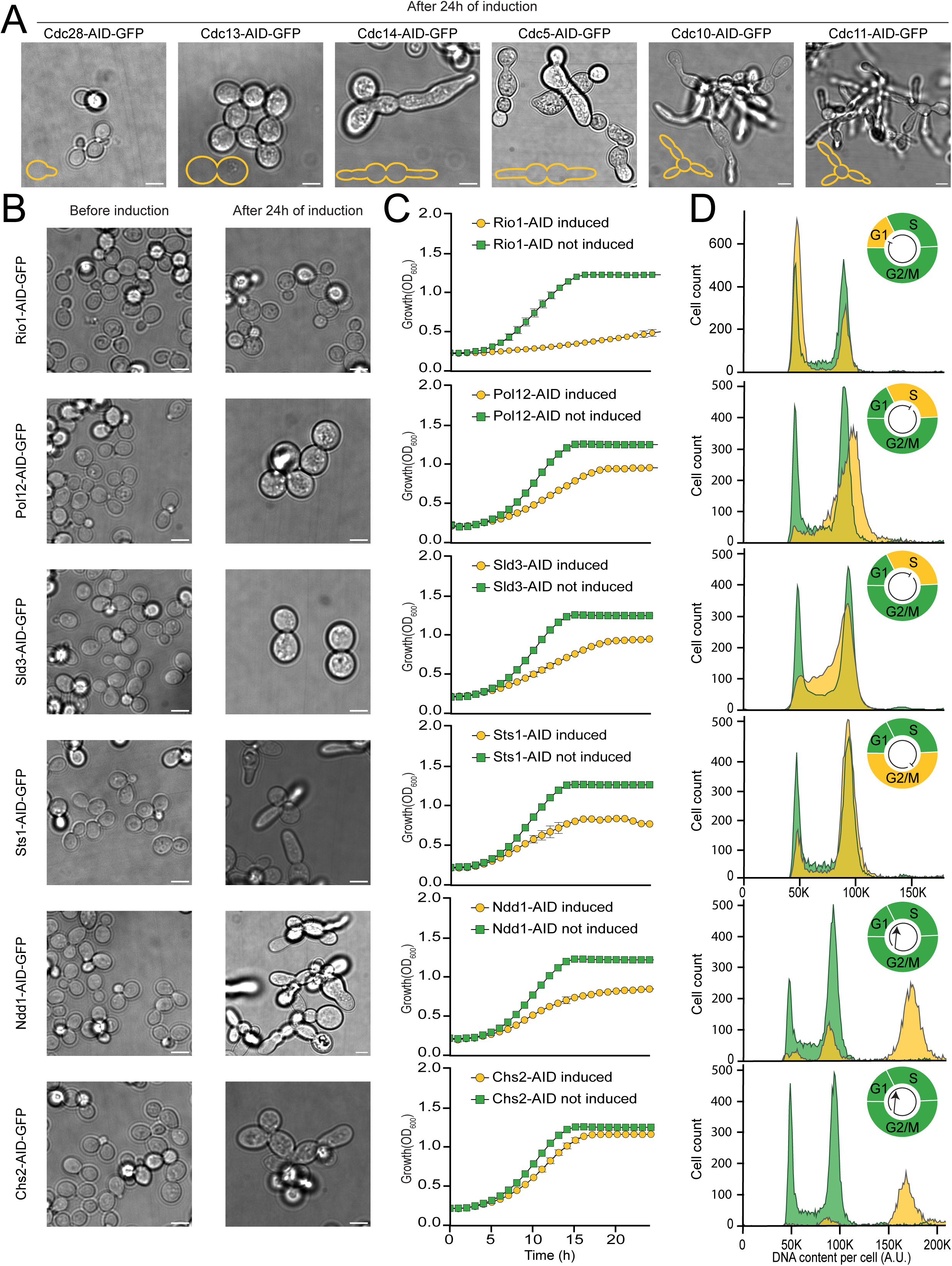
The C’AID-GFP library facilitates the study of CDC genes. A) Brightfield microscopy images of strains previously assigned as CDCs, after induction of the degron system for 24h. A small schematic, recreating the original cartoons (Hartwell et al. 1973) is depicted in the lower left corner of each image. Scale bar 5µm. B) Brightfield microscopy images of strains here suggested as CDCs, before and after induction of the degron system for 24h. In every case, a distinct and synchronized altered morphology can be observed after protein depletion. Scale bar 5µm. C) Growth curves for strains in B, with induction of the degron (orange curve) or not (green curve). Growth defects or lethality are seen upon depletion of the indicated tagged protein. D) Flow cytometry measurements for strains in A, with or without a 3h induction. Without induction (green), a typical histogram for cycling cells is appreciated in each case, while different phenotypes point to issues in the cell division cycle in each protein-depleted case (orange). A schematic of the cell cycle (upper right corner of each panel) shows which phase is affected (orange blocks and arrows).

Encouraged by the sensitivity of our screen, we decided to follow up on some of the strains not previously annotated as CDCs. In addition to the synchronized altered morphology (Figure 5B), the induction of degradation led to growth defects (Figure 5C) and altered flow cytometry profiles (Figure 5D). Notably, we observed cases of cell cycle arrest in various phases, some with literature support: G1 (e.g., Rio1 (Angermayr, Roidl, and Bandlow 2002; Iacovella et al. 2015)), G2 (e.g., Npl4 (Hsieh and Chen 2011) and Sts1), as well as issues in S phase (e.g., Pol12 (Yu et al. 2006) and Sld3). We also documented instances where cells continued to duplicate their DNA without undergoing cellular division (e.g., Chs2 (Schmidt et al. 2002) and Ndd1 (Loy, Lydall, and Surana 1999)). These results collectively pinpoint the utility of the C’ AID-GFP library as a valuable tool for investigating the cell cycle and cell division processes.

## Discussion

Cellular perturbation has been an invaluable approach to uncovering the role of many components in the cell. Inducible systems, in particular, enable the study of dynamic processes and essential components. In this study, we (see also similar library created in parallel (Gameiro et al. 2024)), have created the C’ AID-GFP library, a novel proteome-wide collection designed for on-demand protein depletion to systematically explore cellular processes. Leveraging the versatility of the AID2 system (Yesbolatova et al. 2020) and the efficiency of SWAT libraries (Meurer et al. 2018; Weill et al. 2018; Yofe et al. 2016), we developed a comprehensive tool for investigating protein function and cellular responses.

We demonstrated the robustness of our approach, with near 90% of the expressed proteins responding to induction of the degron (Figure 2A). Moreover, we showcased its applicability by revisiting seminal yeast screens, recapitulating their results, and extending them. We found our system to be highly complementary with other collections that focus on essential genes, such as the YETI-E (Arita et al. 2021) and the TET-off (Mnaimneh et al. 2004) (Supplementary figure 3B), as it effectively evokes the essentiality of lowly expressed proteins. Moreover, our library represents the majority of yeast genes, allowing us to also uncover novel essential genes and new phenotypes for non-essential ones.

Indeed, among our results, we identified essential proteins absent or misannotated in the YKO library (Supplementary figure 3A). Reports in the scientific literature positively support our results with high-confidence demonstrating how the C’ AID-GFP library is a well-suited tool to elicit induction-provoked lethality. In our list of high-confidence new essential proteins, we encounter *YIL102C-A*. Upon closer inspection of this protein, labeled as uncharacterized by the *Saccharomyces* Genome Database (SGD) (Engel et al. 2022; Wong et al. 2023), it appears to have been already characterized as a regulatory subunit of dolichyl phosphate mannose (DPM) synthase (Piłsyk et al. 2020). To this end, we propose *YIL102C-A* to be renamed as *DPM2*, as it’s human counterpart (Maeda et al. 1998).

To ensure widespread accessibility to the scientific community, we openly distribute this library and have compiled our results into yDIMMER (yeast Degron Induced Mitigation for Molecular Exploration of Responses), a user-friendly database: www.ydimmer.com (Supplementary Figure 6). Visualization of any strain present in our library, before or after either short or long induction times is accessible through this platform. Additionally, we provide access to quantitative information about each strain’s relative growth in different media as well as their mitochondrial morphology. The website supports single gene searches, bulk download of information, and filtering by specific properties (e.g., strains with growth defects in respiration media).

In conclusion, our study introduces a powerful tool for functional genomics research, opening up new possibilities for investigating cellular processes. The C’ AID-GFP library, with its ability to systematically deplete proteins and assess cellular responses, promises to accelerate discoveries in yeast biology and beyond.

## Acknowledgement

We thank Noga Preminger and Naama Zung for critical reading of the manuscript. We thank Nir Friedman and Daphna Joseph-Strauss for help with plasmids, we thank Michael Knop for generously sharing the C’ SWAT library and donor plasmids. We thank Emmanuel Levy for reagents and Naama Zung for the Upc2-GFP strain. We are thankful for the amazing work of our support team: Hadar Meyer, Hanni Naor and Reut Ester Avraham; the sequencing unit team and Amir Szitenberg from the Mantoux Bioinformatics G-INCPM unit, at the Weizmann Institute for their service with the library sequencing; and Ronen Hayun, Eli Hotoveli, Arie Pinto and Maayan Maron from the Internet and Mobile Section for their service developing the yDIMMER website and Assaf Glik and Lior Michael from the server unit for providing safe storage and access to the yeast cell images. This study was supported by the Minerva foundation and a Chan Zuckerberg Initiative (CZI) grant (2023-331952). This research was also supported by the Institute for Environmental Sustainability (IES) and the Knell Family Center for Microbiology at the Weizmann Institute of Science. R Valenti was supported by the Alvin, Myra, and David Kaye memorial award for excellence in Life Sciences related research for international students. The robotic system of the Schuldiner lab was purchased through the kind support of the Blythe Brenden-Mann Foundation. MS is an Incumbent of the Dr. Gilbert Omenn and Martha Darling Professorial Chair in Molecular Genetics.

## Materials and Methods

### Yeast strains and plasmids

All yeast strains used in this study are listed in Supplementary Table 2. Strains were constructed either by transformation, using the lithium acetate-based transformation protocol (Daniel Gietz and Woods 2002), or by SWATting (Meurer et al. 2018; Weill et al. 2018; Yofe et al. 2016). All plasmids used are listed in Supplementary Table 3. The donor plasmid was constructed from a plasmid kindly provided by Nir Friedsman lab and the original SWAT donor plasmid. Primers were designed with the Primers-4-Yeast web tool (https://www.weizmann.ac.il/Primers-4-Yeast/ (Yofe and Schuldiner 2014)), Supplementary Table 4, including all the check primers to verify the library clones used in the figures. The SWAT donor strain was constructed by integrating TIR1(F74G) into the ho locus of the original SWAT donor strain (yMS2085; (Weill et al. 2018)). The resulting strain was transformed with a SWAT donor plasmid encoding the C’ AID-GFP tag and crossed with the C’ SWAT library (Meurer et al. 2018).

### Yeast growth

Yeast cells were grown on solid media containing 2.2% [w/v] agar (Formedium #AGA0X) or liquid media; at 30°C. Antibiotics nourseothricin (NAT, WERNER BioAgents “ClonNat” #5.XXX.000) at 0.2 g/l and geneticin (G418, Formedium #G418X) at 0.5 g/l were used for library maintenance and overnight pre-cultures. Unless stated otherwise, the media used was YPD (YPD broth 5% [w/v] Formedium #CCM02XX: 2% [w/v] peptone, 1% [w/v] yeast extract, 2% [w/v] glucose). For microscopy, the media used was synthetic minimal media (SD; 0.67% [w/v] yeast nitrogen base (YNB) without amino acids and with ammonium sulphate (Formedium #CYN04XX), 2% [w/v] glucose), supplemented amino acid OMM mix (Hanscho et al. 2012) (SD complete). For the essentiality test in minimal media, the same SD was used, but only supplement with histidine 3.5mg/L (Formedium #DOC0142), methionine 4mg/L (Formedium #DOC0166, and uracil 4mg/L (Formedium #DOC0212), required for the survival of our strains given their genetic background (SD minimal). For the essentiality test in respiration media we used YPGlycerol, prepared in the same manner as YPD but with 3% [w/v] glycerol (CAS56-81-5; J.T. Baker: Product code: 15557974) instead of glucose. In all cases, 5-Ph-IAA (Inducer for Auxin Inducible Degron 2 System; CAS: 168649-23-8; Bio Academia: Product code: 30-003-10) was used at a working concentration of 5µM and added to the media for the time indicated in each figure. A stock solution (100mM in DMSO/20.000X) was kept in aliquots at -20°C and only thawed/frozen for a few cycles. For solid media, the 5-Ph-IAA was incorporated together with the antibiotics, when the temperature of the media was enough to keep the agar liquid, but low to prevent damaging the auxin. Utilizing a working concentration of 1µM provided the same results as 5µM in time course analysis, data not shown.

### Yeast library generation

The C’ AID-GFP SWAT library generation was performed as described (Weill et al. 2018). In short, a RoToR array pinning robot (Singer Instruments) was used to mate the donor strain (Table S4) with the parental C′ SWAT library. The mating was followed by sporulation and selection to generate the desired haploid α library, harboring all the required elements (Cohen and Schuldiner 2011; Tong and Boone 2007): HO::NAT-TEF2p-OsTIR1(F74G), XXX(ORF)-AID*-eGFP::G418. For SceI-mediated tag swapping, the library was grown on YPGalactose (2% [w/v] peptone, 1% [w/v] yeast extract, 2% [w/v] galactose) and then moved to SD containing 5-fluoroorotic acid (5-FOA, Formedium) at 1 g/l. Finally, NAT and G418 were employed to select the strains that successfully completed the SWAT procedure. Images of the finished library were taken with a CCD camera (Amersham ImageQuant 800 systems, Cytiva). Images were analyzed with SGA tools (Wagih et al. 2013) to extract the colony sizes and compared with the images of the parental library (C’ SWAT) to determine colony survival.

### Library validation by targeted sequencing (Anchor-Seq)

The Anchor-Seq method (Meurer et al. 2018) for targeted sequencing of the pooled library was utilized. In brief, yeast strains from a grown agar plate of the library were collected, washed with buffer, aliquoted (100-150 mg per tube), and stored at -20°C until use. The Zymo Research YeaStar kit (cat. No.: D2002) was used as per the manufacturer’s direction to extract genomic DNA from one aliquot per plate, with a typical yield of 115-125 ng/uL in 60 uL. To fragment the DNA, the NEBNext UltraTM II FS DNA Library Prep Kit for Illumina® (Cat. No.: 7805S) was used with fragmentation lasting for 5 minutes at 37°C. The bubble-shaped adaptor was achieved by resuspending oligos #1 and #2 (Supplementary Table 5) in TE buffer and annealing them in annealing buffer via heat denaturation and slow cooling. The fragmented DNA received 2.5 uL of the bubble-shaped adaptor at a concentration of 15 mM that was ligated for 15min at 20° C in a LAB cycler (a SensoQuest PCR model). The NEBNext Sample Purification SPRI Beads were used for size selection and cleanup of adaptor-ligated DNA. A selective PCR preformed with oligos #3 and #4 was performed for 18 cycles, followed by a new round of beads purification and a second PCR with primers #5 and #6, for 15 cycles. A final size selection and cleanup were performed before DNA quantification with a Qubit Flex Fluorometer (Thermo Fischer Cat. No. Q33327) and normalization to 8 nM per tube. Equal volumes from the four normalized samples were pooled together and spiked with 25% Phi X DNA before being subjected to NGS sequencing. 15. NGS. We used Illumina MiSeq, v3_600 cycles flow cell kit (Cat. No. MS-102-3003), with a paired-end protocol allocating 310 read cycles from the Rd1 and Rd2 directions and 6 bases read for the Index 1 barcodes. The barcodes assigned to each sample (refer to Supplementary Table 5) were utilized to match the reads with their respective 1536 plate. Sequencing and Bioinformatic analyses were performed at the Mantoux Bioinformatics G-INCPM, the Weizmann Institute. FASTQ files accession number: PRJNA1121868.

### Automated high-throughput fluorescence microscopy

For the short-term microscopy screen, the library was transferred from agar plates into 384-well plates for growth in 100µl YPD media with antibiotics overnight (ON) at 30°C using the RoToR arrayer The ON culture was back diluted into 384-well plates in YPD to an OD_600_ of ∼0.2. After 4h, 50µl from each well were transferred to a glass-bottom 384-well microscopy plate (Matrical Bioscience) coated with concanavalin A (Sigma-Aldrich). The logarithmic phase cultures were allowed to bind to the bottom of the plate for 20 minutes. After incubation, wells were washed twice with SD complete media to remove non-adherent cells. The plates were then transferred to an Olympus automated inverted fluorescent microscope system using a robotic swap arm (Hamilton). Cells were imaged at room temperature (RT) in SD complete media using an Olympus UPLFLN 60X air lens (NA 0.9), a CSU-W1 Confocal Scanner Unit (Yokogawa), equipped with a 50µm pinhole disk, and with an ORCA-flash 4.0 digital camera (Hamamatsu), using the ScanR software (version 3.2 Images were recorded with 488 nm laser illumination for the GFP channel (excitation 488nm, emission filter 525/50 nm), with 500ms exposure and 70% laser intensity, mild conditions to minimize photobleaching (compared to long-term screen below). Bright-field images were also taken, with 200ms exposure and 100% halogen bulb. Two positions were imaged per well and the software autofocus was used to ensure the cells were imaged in their central plane for proper comparison. After imaging, the plates were removed from the microscope, 5-Ph-IAA was added to a final concentration of 5µM per well and the plate was incubated for 30 min at RT. After incubation, the plate was re-imaged as previously. The same positions were re-visited. All the steps in the automated imaging process were done by an automated imaging platform EVO freedom liquid handler (TECAN).

For the long-term microscopy screen, the procedure was as described above, with the changes here noted. The back dilution culture (100µl) was split into two: 50µl were transferred to the microscopy plate and imaged (before induction), while the remaining 50µl received 5-Ph-IAA to a final concentration of 5µM. These plates were grown for 20h at 30°C to become the ON plate for the induced cultures. After 20h, 5µl were back diluted into 95µl of SD complete with 5µM 5-Ph-IAA. After 4h of back dilution, 50µl from each well were transferred to the microscopy plate. Cells were allowed to bind to the bottom of the plate for 15min. After the incubation, the media was replaced by fresh media, with 50nM Mitotracker™Orange CMTMRos (ThermoFisher Scientific) and 5µM 5-Ph-IAA. The dye was incubated for 15 min and washed with SD complete with 5µM 5-Ph-IAA. Cells were imaged as described previously, in SD complete with 5µM 5-Ph-IAA (24h induced cells). Since the cells were not re-imaged, harsher illumination conditions were used to improve GFP visualization (1000ms exposure time, 80% laser intensity). In the induced case, Mitotracker™Orange was imaged with an exposure of 600ms, 60% laser intensity (excitation 561nm, emission filter 617/73 nm).

### Low-throughput fluorescence microscopy

Images displayed in Figure 2 and 4 and in Supplementary Figure 3 and 4 were acquired in the same setup as before, but with an Olympus UPLFLN 100X oil objective (NA 1.3).

Images displayed in Figure 1 were acquired at RT using a VisiScope Confocal Cell Explorer system, composed of a Yokogawa spinning disk scanning unit (CSU-W1) coupled to an inverted Olympus IX83 microscope with a 60X oil objective (NA 1.4). The excitation wavelength was 488nm, with an exposure time of 600ms and 80% laser intensity). Images were taken by a PCO-Edge sCMOS camera controlled by VisiView software (Visitron Systems version 3.2.0).

The microscopy images were cropped and colored in ImageJ (Schneider, Rasband, and Eliceiri 2012). The brightness and contrast of images were linearly adjusted so that all GFP images of the same strain (before and after induction) have the same parameters. For MitoTracker images, all the images in the same panel, irrespective of the strain or induction state, share the same parameters. In cases where the cell boundaries were not visible, the Dotted line plugin for ImageJ was used to denote the cell edge.

### Essentiality assays

The C’ AID-GFP library was replicated in 1536 format in agar with antibiotics, using the RoToR arrayer (Singer). Two copies were replicated in each media tested (YPD, SD minimal and YPGlycerol, see yeast growth) with 5µM 5-Ph-IAA and two replicas without. After 24h of incubation at 30°C (or 48h for YPGlycerol), cells were refreshed into new plates using the RoToR arrayer. After a second period of incubation (24h at 30°C, 48h for YPGlycerol), all the plates were imaged with a CCD camera (Amersham ImageQuant 800 systems, Cytiva). Images were analyzed with SGA tools (Wagih et al. 2013) to extract the colony sizes. An average of both replicates was used to calculate the ratio between the colonies grown with 5-Ph-IAA and the colonies grown without (not induced). As controls, a WT strain (YMS3553, Supplementary Table 2) and three library strains known to give induction-provoked lethality were incorporated into each plate in triplicates. Since our metric was based on a ratio between induced and uninduced strains, positional effects (such as better growth in the plate edges) are expected to cancel out. However, we normalized the scores of each plate and each media to facilitate comparisons using the median value per plate, after removal of outliers.

### Growth assays

Cells were grown ON in YPD with antibiotics and the culture was washed from old media and back diluted to OD_600_ 0.2 in fresh YPD containing or not 5µM 5-Ph-IAA. Cells were incubated at 30°C with agitation and growth was monitored every hour for 24h with a Spark multimode microplate reader (TECAN) by measuring the OD_600_. In the cases of the GAL promotor swap, the ON growth in YPD served to deplete the mRNA from the cells. The cells were then grown on YPD (repressive conditions) or YPGalactose (prepared as YPD but with 3% galactose). At least three replicates were used in every case.

### Responsiveness under different conditions

Protein depletion was successfully verified under different media conditions, after 45 minutes of induction, including: SD (MSG) complete (prepared as SD complete but 0.1% MSG (Minerva); 0.17% YNB without amino acids and without ammonium sulfate (Minerva); SD (MSG) complete with 5µg/ml Terbinafine (Minerva) or 2mM DTT (Sigma); SD (MSG) complete without thiamine (Minerva); without biotin (Minerva); without copper (Minerva); without iron (Minerva) or without phosphate (Minerva); SGalactose (MSG) complete; DDW and conditioned media SD (MSG) complete after 48h of yeast growth, followed by sterilization by addition of antibiotics.

### Creation of grid images of individual yeast cells

To maximize the information content of each microscopy image, single yeast cells were automatically segmented from the brightfield image using a neural network within ScanR Analysis software. Detected cells were individually cropped by delimitating a square bounding box of 65×65 pixels centered on them. A montage image assembled up to 400 individual cells from the same strain laying them out in a grid of 20 by 20 cells.

Thus, for each strain, we generated a cell grid image of dimension 1300×1300 pixels for each combination of conditions tested (short vs. long induction and before vs. after induction). In addition, cell masks were transferred from brightfield (“TransCon”) across the fluorescent channels (488nm and 561nm) to yield one cell grid montage per channel. To facilitate visualization, the contrast of each cell-grid image was adjusted such that 0.35% of pixels were saturated. Importantly, we preserved fluorescence intensities across cells, thus allowing us to assess for noisy protein expression (e.g. high variance across cells) or intensity variations between timepoints (e.g. before or after 30mn/24hr induction). To save storage space and increase performance in loading images, intensities of brightfield images were scaled down to 8 bits since intensity variations were not useful. Custom python scripts to generate cell grids are available in https://github.com/benjamin-elusers/ydimmer.

### Analysis displayed in images

#### “SWATting” survival

Images of the C’ SWAT before library creation and of C’ AID-GFP in agar plates were quantified with SGA tools (Wagih et al. 2013), to estimate the colony “SWATting” survival. A colony was considered as present if the quantification was over 50 A.U. Supplementary Figure 1A.

#### “SWATting” efficiency

The top 500 most abundant proteins (Ho et al. 2018), whose strain were present in the library, were evaluated for fluorescence. Since these highly abundant proteins should provide a clear signal, the percentage of strains with signal detected can serve as a proxy for “SWATting” efficiency, meaning the genetic modification with the AID tag and GFP module were successful. Supplementary Figure 1B.

#### Sequencing validation and coverage

From the sequencing output (see Library validation by targeted sequencing (Anchor-Seq)) we classified the strains into categorical results. The categories represent whether the strain was validated by sequencing, not found, or found with an undesired modification (mutation). The percentage of correct sequences over the total found gave us the sequencing validation, Supplementary Figure 1C; while the percentage recovered over the submitted gave us the coverage, Supplementary Figure 1D.

#### Fluorescence and responsiveness classification

Images acquired in the long and short induction screen were processed with ScanR Analysis, to segment individual cells and measure the mean green fluorescence per cell. For each strain, cells with signals outside of a range of two standard deviations (SD) were removed and the geometric mean, SD and number of cells (n) were reported, Supplementary Table 1. Strains where the fluorescence is bigger than the brightest non-fluorescent control were analyzed in a quantitative manner to determine responsive (significant GFP signal reduction after induction). For the remaining strains, mainly organellar ones, visual examination was used to determine if the GFP signal was present and if it was reduced upon induction. Values and individual results for each strain are listed in Supplementary Table 1, while the aggregate of the annotations was utilized to calculate the percentages displayed in Figure 2A. The individual values for fluorescent strains were plotted in Figure 2B (long induction) and Supplementary Figure 2 B (short induction) in RStudio (Posit team 2023) with the ggplot2 package (Wickham 2016).

#### Responsiveness across cellular compartments

The sub-cellular protein localization for each fluorescent strain in the library was adapted from (Huh et al. 2003) to include only the first category listed and only the categories listed in Figure 2D. For that, “early Golgi” and “late Golgi” were considered together with “Golgi”; “nucleolus” was integrated to “nucleus”; and “actin”, “endosome”, “ER to Golgi”, “microtubule” and “spindle pole” were considered under “punctate composite”. The percentage of responsive strains for each category is displayed in Figure 2D.

For mitochondrial sub-compartments, the annotations from (Williams et al. 2014) were taken and used to compute the percentage responsiveness across mitochondrial compartments, Figure 2E, and further considering co-vs post-translational translocation, Supplementary Figure 2C.

For bias in responsiveness according to orientation, the localization of the C’ of the ER proteins(as determined by (Huh et al. 2003)) was computed by the majority rule from the algorithms listed in Topology yeast (Weill et al. 2019). Annotations are listed in Supplementary Table 1, and group percentage of responsiveness are displayed in Supplementary Figure 2D.

#### Fluorescence and responsiveness across protein abundance

All strains present in the library were segregated by protein abundance (Ho et al. 2018) into deciles, with decile 1 clustering the most abundant proteins and decile 10 the least. The percentage of the strains of each decile with detected fluorescence or responsiveness are presented in Supplementary Figure 2E and F, respectively.

#### Go slim analysis for proteins essential under different conditions

The list of strains with relative growth below 0.5 in induced vs uninduced conditions (hits) in each media were taken to generate the Venn diagram displayed in Figure 2C. All the GO slim terms (Ashburner et al. 2000; The Gene Ontology Consortium et al. 2023) associated with each of the hits were grouped (i.e. minimal media includes 480 hits). Since most hits were shared (347 strains were this in every condition tested), to select the top three GO slim terms associated with each media, the exclusive hits were taken (meaning the strains that were only found to be essential in that specific condition, for minimal media, 73 out of 480 hits). For reporting the fraction of each GO slim term that was recovered, all the strains associated with that specific GO term were now utilized (i.e. “Amino acid metabolic process”, GO:0006520, was the term strongly associated with the hits exclusively in minimal media (22/73); yet, if we consider all the 480 hits in minimal media, 27 hits are present). The percentage of the hole GO slim term annotations affected is reported, regardless of whether those strains were present or not in the library (continuing the example, “Amino acid metabolic process” has 151 genes associated to it, but 137 of those are represented in the C’ AID-GFP library, and 77 have been certified as responsive from fluorescent analysis; this gives a 17.9% of the total annotations for the term (27/151) as hits). The top three GO terms for each category are listed under the titles for each media in Figure 2C, with the percentage of each GO slim term recovered as hits in the screen. For the hits shared in all the conditions, only the 347 strains were considering when quantifying percentage of GO term recovery.

#### Comparison with other systems

To compare the list of strains classified as essential in the YKO (Giaever et al. 2002; Winzeler et al. 1999) with the C’ AID-GFP results, the relative growth in rich media in induction vs not induction was considered. Venn diagrams displaying the overlap between the 560 essential genes defined in the YKO and defined as responsive in the C’ AID-GFP library and the strains with a relative growth below 0.5 (permissive threshold) or 0.1 (stringent threshold) are displayed in Supplementary Figure 3A.

To compare the C’ AID-GFP library with other inducible systems, the list of proteins classified as essential in the YKO (Giaever et al. 2002; Winzeler et al. 1999) and present in the C’ AID-GFP library (irrespective of whether the strain is responsive), 650 proteins, was separated into quintiles, each with 130 proteins, by their mRNA abundance (Lipson et al. 2009), with quintile 1 representing the most abundant ones. Qualitative classification of the strains from the TET-off (Mnaimneh et al. 2004) and the YETI-E (Arita et al. 2021) collection into “responsive”, “non-responsive” and “absent” allowed for the generation of the scatter plots in Supplementary Figure 3B of percentage genes with affected phenotypes vs mRNA abundance.

#### Mitochondrial signal quantification and determination of MDM-like phenotypes

The images acquired of the MitoTracker die after 24h of induction (see Automated high-throughput fluorescence microscopy, long term) were processed as in Fluorescence and responsiveness classification, with a further segmentation of the mitochondria per cell done with the edge segmentation tool in ScanR Analysis. Changes in fluorescence intensity, mitochondria number and relative growth in respiration media were instrumental in determining a strain as displaying MDM-like phenotypes, together with a blind manual examination of the images obtained, Supplementary Table 1. The MitoTracker signal of each strain was ranked to generate the plot in Figure 4B.

#### Ergosterol biosynthesis pathway proteins

Strains harboring the degron tag in a protein related to the ergosterol biosynthesis pathway were induced or not for 4h and then imaged as described in low throughput microscopy. The quantitation of GFP and MitoTracker fluorescence was done as detailed in fluorescence and responsiveness classification and displayer together with an schematic of the pathway adapted from (Jordá and Puig 2020) in Supplementary Figure 4C.

#### Synchronized and altered morphology

Bright-field images acquired after 24h of induction (see Automated high-throughput fluorescence microscopy, long term) were manually examined to determine candidates displaying synchronized and/or altered morphology, Supplementary Table 1.

#### Western blot

Cells were grown ON in YPD with selections and used to inoculate 125ml of YPD to OD_600_ 0.05. Cells were incubated with shaking at 30°C until OD_600_ 0.2, where they were spit into 18 identical cultures. The first series of technical replicates received 5µM 5-Ph-IAA (time point 180 min), and the subsequent groups received it after 1.5h, 2h, 2.5h and 2.8h so that after 3h of growth all the cells were ready for harvesting. Cells were harvested by centrifugation, followed by flash freeze in liquid nitrogen. Protein extraction, SDS-PAGE, and western blotting were preform as described previously (Eisenberg-Bord et al. 2021). Briefly, the cells were lysed with 8M urea-based lysis buffer with protease inhibitors and glass beads-beating (Scientific industries). Lysates were denatured by the addition of SDS (final concentration ∼2%) and a 45°C incubation for 15min. Denatured lysates were centrifuged to separate cell debris. Loading buffer containing DTT (final concentration ∼25mM) and incubated at 45°C for 15min. 30μg sample was loaded onto 12% agarose gels and separated with electrophoresis, then transferred onto nitrocellulose membrane using the Trans-Blot Turbo transfer system (Bio-Rad). Membranes were blocked in SEA BLOCK buffer (Thermo Scientific), incubated with primary antibodies (anti-GFP, Abcam, ab290 1:1000 and Anti-actin, Abcam, ab170325 1:5000), washed, and incubated with fluorescent secondary antibodies for 1h (Li-COR, 926-32210, 1:10000 and Abcam, ab216777, 1:10000). After washing, the membranes were imaged on the LI-COR Odyssey Infrared Scanner.

#### Flow cytometry

The strains were grown ON at 30°C and shaking, 180 RPM (New Brunswick model Innova 44) in Falcon 15 ml rounded-bottom tubes (Corning) with a loose a loose cap with 1 ml of S.D. complete [MDRV1] medium supplemented with Nat antibiotics. The O.N. cultures were back-diluted into 2ml of the same media to an OD600 of 0.2. The cultures were allowed to grow until reaching OD600 of 0.5, when they split in half. In one-half of each sample, 5µM 5-Ph-IAA was added before incubation was continued for another three hours and then stopped by adding 2ml of Ethanol (final concentration 66%). After the Ethanol addition, the cultures’ tubes were vortex-mixed rigorously before being stored at -20°C until further use (a minimum of ON). The cell cultures were prepared for DNA visualization by flow cytometry analysis as previously described (Rosebrock 2017). In short, following ethanol fixation, cells were treated with RNAse A (Sigma) to remove RNA and, after that, with Proteinase K (Sigma) to remove proteins. The treated cell samples were then stained with 2.5µM Sytox Green (Thermo Fischer) to visualize DNA content before analyzing the samples at the Flow Cytometry Core Facility of the Weizmann of Science [MDRV2] [MDRV3] using the BD FACSAria™ III Cell Sorter equipped with a 100 nm nozzle and controlled by BD FACS Diva software v8.0.1 (B.D. Biosciences). DNA content of the cell population was collected by excitation of the Sytox Green at 488 nm and monitoring its emission at 502 with long-pass and 530/30 bandpass filters and plotted as the area of the Sytox-green signal Vs. cell count. Further analyses were performed using FlowJo software v10.2 (Tree Star).

**Supplementary Figure 1. The C’ AID-GFP library extends the degron approach to a proteome level.**

A) Pie chart displaying the rate of survival of strains following the SWATting procedure. Colony presence was measured for the parental (C’ SWAT library) and derived (C’ AID-GFP) libraries, to calculate the 91% survival rate. B) Pie chart displaying the fluorescent status of the top 500 most expressed proteins in the cell (Nash et al. 2020). For such abundant proteins a fluorescent signal was expected, therefore the estimation of the SWATting efficiency corresponds to the 94% strains with signal. C) Pie chart displaying the proportion of functional tags detected by sequencing in the library. Of 3308 strains detected by pooled sequencing, only 12 displayed indels that would render the tag dysfunctional. D) Pie chart displaying the sequencing coverage. Of 3670 colonies sequenced, 90% were successfully recovered, displaying a functional tag.

**Supplementary Figure 2. Characterization of responsiveness across different parameters highlights the factors influencing the C’ AID-GFP efficiency.**

A) Fluorescent images of selected proteins imaged after no (0 min), short (30 min) or long (24h) periods of induction. Different strains respond to the treatment with different kinetics, with some being depleted after short induction times (Lia1), while others require long induction times (Mrh1). In cases where the signal is too dim, the outline of the cells is depicted with a dotted line. Scale bar 5µm. B) Dot plot showing the mean GFP fluorescence intensity per cell for each strain before or after a short induction time. The background autofluorescence is marked in dotted lines (A.F., mean and two standard deviations in each direction). The diagonal line indicates the expected location for irresponsive strains. Already in this short induction times there is a clear response in many strains. C and D) Schematics depicting the differences in responsiveness for proteins that localize to mitochondria (C) and are co-vs post-translationally translocated (Williams et al. 2014), or to the Endoplasmic Reticulum (D) and have their C’ exposed in the cytosol or hidden in the lumen. Proteins targeting to mitochondria only after their synthesis is complete (post translational) are more likely to be depleted upon induction of the AID system. Proteins with the C’ exposed to the cytosol are also more likely to be degraded than their counterparts with hidden C’. This reflects differences in the accessibility of the AID tag to the TIR1 adaptor protein, the ubiquitination machinery and the proteasome, required for its induced depletion. Translocation and topology information were taken from the literature (Jan, Williams, and Weissman 2014; Weill et al. 2019). E) Bar plot of percentage of fluorescent strains vs abundance decile, with decile 1 being the most abundant proteins (Nash et al. 2020). As expected, abundant proteins are more likely to have a fluorescent signal. F) Bar plot of percentage of responsive strains vs abundance decile (Nash et al. 2020). For the fluorescent strains in each decile, no clear correlation between abundance and responsiveness is detected. Responsiveness remains above 80% for all the deciles.

**Supplementary Figure 3. The C’ AID-GFP library is complementary to other approaches.**

A) Venn diagrams comparing the results from the C’ AID-GFP library with the YKO collection (Giaever et al. 2002) for hits defined with a permissive (<0.5 relative growth) or stringent (<0.1 relative growth) thresholds. B) Graphs showing the comparison of the C’ AID-GFP library vs other inducible collections such as the YETI-E (Arita et al. 2021) and TET-off collection (Mnaimneh et al. 2004) across protein abundance, where quintile 1 groups the proteins with the most abundant mRNA and 5 the least (Lipson et al. 2009). C) Fluorescent images demonstrating that the AID system works robustly under different conditions. Scale bar 5µm.

**Supplementary Figure 4. Known proteins and pathways required for proper mitochondrial morphology and distribution display altered mitochondrial phenotypes as expected.**

A) Fluorescent images of known MDM strains imaged without induction or after 24h of induction. The mitochondria morphology (MitoTracker channel) is severely affected after protein depletion (GFP channel). Scale bar 5µm. B) Fluorescent images for the proteins related to the ergosterol biosynthesis pathway before and after 4h of induction. Mitochondria with brighter signal (quantification od mean MitoTracker signal per cell +/-SD in the MitoTracker images) and with altered morphology (magenta) can be seen for most proteins C) Schematics of the ergosterol biosynthesis pathway with quantification of the strains shown in B for the MitoTracker signal (purple no induction, magenta 4h induction) and GFP (green no induction, gray 4h of induction). Dashed vertical lines indicate the background fluorescence in the control. Changes are observed irrespective of whether the proteins affected participate in the pre (blue and green) or post (orange) squalene module of the pathway. Image adapted from (Jordá and Puig 2020).

**Supplementary Figure 5. A Cdc48 complex member displays CDC characteristics.**

A) Left: Brightfield microscopy images of Cdc48, before and after induction of the degron system for 24h. A distinct and synchronized altered morphology can be observed after protein depletion as previously shown. Scale bar 5µm. Center: Growth curves of Cdc48, with induction of the degron (gray curve) or not (green curve). Lethality is seen upon depletion of Cdc48. Right: Flow cytometry measurements for Cdc48, with or without 4h induction. Without induction (green), a typical histogram for cycling cells is appreciated, while an arrest in G2/M stage can be seen upon protein depletion (gray and upper right corner schematic). B) Same as A, but for Npl4, a Cdc48 interactor. The growth curve is identical to the one displayed in Figure 3C.

**Supplementary Figure 6. yDIMMER database allows access to the C’ AID-GFP strains images and information.**

A) Example entry for the protein Cys3. Left: Information about the gene, including external resources as the link to the SGD (Engel et al. 2022; Wong et al. 2023), the GPF localization (Huh et al. 2003) and the YKO phenotype (Giaever et al. 2002); as well as the results from the morphology and essentiality analysis conducted with the C’ AID-GFP library. Center: Fluorescent images from the short induction assay, together with their quantification. Right: Fluorescent images form the long induction assay, together with their quantification. Note that the long induction assay was performed under harsher illumination conditions so the quantification of the GFP signal will always be bigger than for the short assay at time 0 min. For the 24h time point there is information from the MitoTracker signal. B) Same entry from the website, but with this time showcasing the zooming options (center) and the option to visualize the bright-field or the MitoTracker signal (for 24h induction).

